# Genome Scale Epigenetic Profiling Reveals Five Distinct Subtypes of Colorectal Cancer

**DOI:** 10.1101/397620

**Authors:** Lochlan Fennell, Troy Dumenil, Gunter Hartel, Katia Nones, Catherine Bond, Diane McKeone, Lisa Bowdler, Grant Montgomery, Leesa Wockner, Kerenaftali Klein, Isabell Hoffmann, Ann-Marie Patch, Stephen Kazakoff, John Pearson, Nicola Waddell, Pratyaksha Wirapati, Paul Lochhead, Yu Imamura, Shuji Ogino, Renfu Shao, Sabine Tejpar, Barbara Leggett, Vicki Whitehall

## Abstract

**BACKGROUND:** Colorectal cancer is an epigenetically heterogeneous disease, however the extent and spectrum of the CpG Island Methylator Phenotype (CIMP) is not clear.

**RESULTS:** An unselected cohort of 216 colorectal cancers clustered into five clinically and molecularly distinct subgroups using Illumina 450K DNA methylation arrays. CIMP-High cancers were most frequent in the proximal colons of female patients. These dichotomised into CIMP-H1 and CIMP-H2 based on methylation profile which was supported by over representation of *BRAF* (74%, P<0.0001) or *KRAS* (55%, P<0.0001) mutation, respectively. Congruent with increasing methylation, there was a stepwise increase in patient age from 62 years in the CIMP-Negative subgroup to 75 years in the CIMP-H1 subgroup (P<0.0001). There was a striking association between PRC2-marked loci and those subjected to significant gene body methylation in CIMP-type cancers (P<1.6xl0^78^). We identified oncogenes susceptible to gene body methylation and Wnt pathway antagonists resistant to gene body methylation. CIMP cluster specific mutations were observed for genes involved in chromatin remodelling, such as in the SWI/SNF and NuRD complexes, suggesting synthetic lethality.

**CONCLUSION:** There are five clinically and molecularly distinct subgroups of colorectal cancer based on genome wide epigenetic profiling. These analyses highlighted an unidentified role for gene body methylation in progression of serrated neoplasia. Subgroup-specific mutation of distinct epigenetic regulator genes revealed potentially druggable vulnerabilities for these cancers, which may provide novel precision medicine approaches.

## Background

Colorectal cancer is a heterogeneous disease characterized by distinct genetic and epigenetic changes that drive proliferative activity and inhibit apoptosis. The conventional pathway to colorectal cancer is distinguished by *APC* mutation and chromosomal instability, and accounts for approximately 75% of sporadic cancers (1, 2). The remaining colorectal cancers arise from serrated polyps and have activating mutations in the *BRAF* proto oncogene, frequent microsatellite instability (MSI), and aberrant genome-wide CpG island (CGI) methylation termed the CpG Island Methylator Phenotype (CIMP) (2, 3).

The development of CIMP is critical in the progression of serrated neoplasia (3). It is well established that CIMP can result in the silencing of key genes important for tumour progression, including the tumour suppressor gene *CDKN2A* and the DNA mismatch repair gene *MLH1* (4, 5). Gene silencing mediated by *MLH1* promoter hypermethylation impairs mismatch repair function which leads to microsatellite instability (5). CIMP can be detected using a standardized marker panel to stratify tumours as CIMP-high, CIMP-low or CIMP-negative (3). Activation of the mitogen-activated protein kinase (MAPK) signaling pathway due to *BRAF* mutation is highly associated with CIMP-high. CIMP-high cancers frequently arise proximal to the splenic flexure and are more common in elderly female patients (2, 3) whilst CIMP-low cancers have been associated with *KRAS* mutation (6, 7).

More recently, consensus molecular subtyping (CMS) was proposed for classifying colorectal cancers based on transcriptional signatures. Guinney and colleagues identified four major molecular subtypes (CMS1-CMS4) (8). CMS1, or MSI immune subtype, is characterized by MSI, *BRAF* mutation and enhanced immunogenicity. CMS2 can be distinguished by chromosomal instability and WNT pathway perturbations. CMS3, or metabolic subtype, is characterized by *KRAS* mutation, CIMP-low status and infrequent copy number alterations. CMS4, or mesenchymal subtype, shows high copy number aberrations, activation of the transforming growth factor-β signaling cascade, stromal infiltration and the worst overall survival. The relationship between CIMP and CMS subtypes is currently unclear.

Methylation is not a phenomenon distinct to neoplasia. Changes in the epigenome also occur with age and in response to environmental factors (9, 10). We have previously shown that the promoter region of certain genes becomes increasingly methylated in normal colonic mucosa with age (9). CIMP-high cancers are identified primarily in older patients (2) hence, age related hypermethylation might prime the intestinal epigenome for serrated neoplasia-type colorectal cancers. Methylation is also critical in the progression of serrated pathway precursors to invasive cancer, primarily through methylation of MLH1 at the transition to dysplasia (11)(12). Thus the natural history of the cancer within the colorectum may dictate the methylation profile of the cancer once malignancy develops.

DNA methylation alone can be insufficient to induce transcriptional repression (13). Gene repression is also associated with repressive histone marks such as the H3K27me3 mark (14), which is catalyzed by the polycomb-repressor-complex 2. Modification of histone tails is catalyzed by a series of enzymes including epigenetic readers, which scan for histone modifications; writers, which effect the addition of a modification; and erasers, which are responsible for the removal of histone marks. Mutations in genes encoding epigenetic enzymes have been shown to occur frequently in cancer (15). Whilst DNA methylation is classically associated with gene silencing, the relationship between DNA methylation and histone modifications has not been fully elucidated, nor has the role of somatic mutations in enzymes that catalyze these epigenetic processes been comprehensively examined.

In this study, we define the extent and spectrum of DNA methylation changes occurring in colorectal cancers and relate this to key clinical and molecular events characteristic of defined pathways of tumour progression. We investigate the role of DNA methylation in the modulation of gene transcription, and assess mutation of genes encoding epigenetic regulatory proteins.

## Results

### Clinical and molecular features of the cohort

Genome wide methylation levels were assessed for a total of 216 unselected colorectal cancers (Table 1). The mean age of patients at surgery was 67.9 years. 29 of 216 (13.4%) of cancers had a *BRAF* V600E mutation, and 75 of 216 (34.7%) cancers were mutated at *KRAS* codons 12 or 13. Mutation of *BRAF* and *KRAS* were mutually exclusive. Patients with *BRAF* mutated cancers were significantly older than patients with *BRAF* wild type cancers (mean age 74.9 vs 66.9, P=0.01). *TP53* was mutated in 78/185 (42.2%) cancers. MSI was significantly associated with *BRAF* mutation (18 of 29 *BRAF* mutant vs 9 of 187 compared with *BRAF* wild-type cancers, P<0.0001). Using the Wiesenberger panel to determine CIMP status(3), 24/216 (11.1%) were CIMP-high, 44/216 (20.4%) were CIMP-low and 148 of 216 (68.5%) were CIMP-negative. CIMP-high was significantly associated with *BRAF* mutation compared with *BRAF* wild-type cancers (19/29 vs 5/186, P<0.0001). CIMP-low was significantly associated with *KRAS* mutation compared with *KRAS* wild-type cancers (26/75, 34.6% vs 18/141, 12.8%, P<0.001).

**Table 1:**
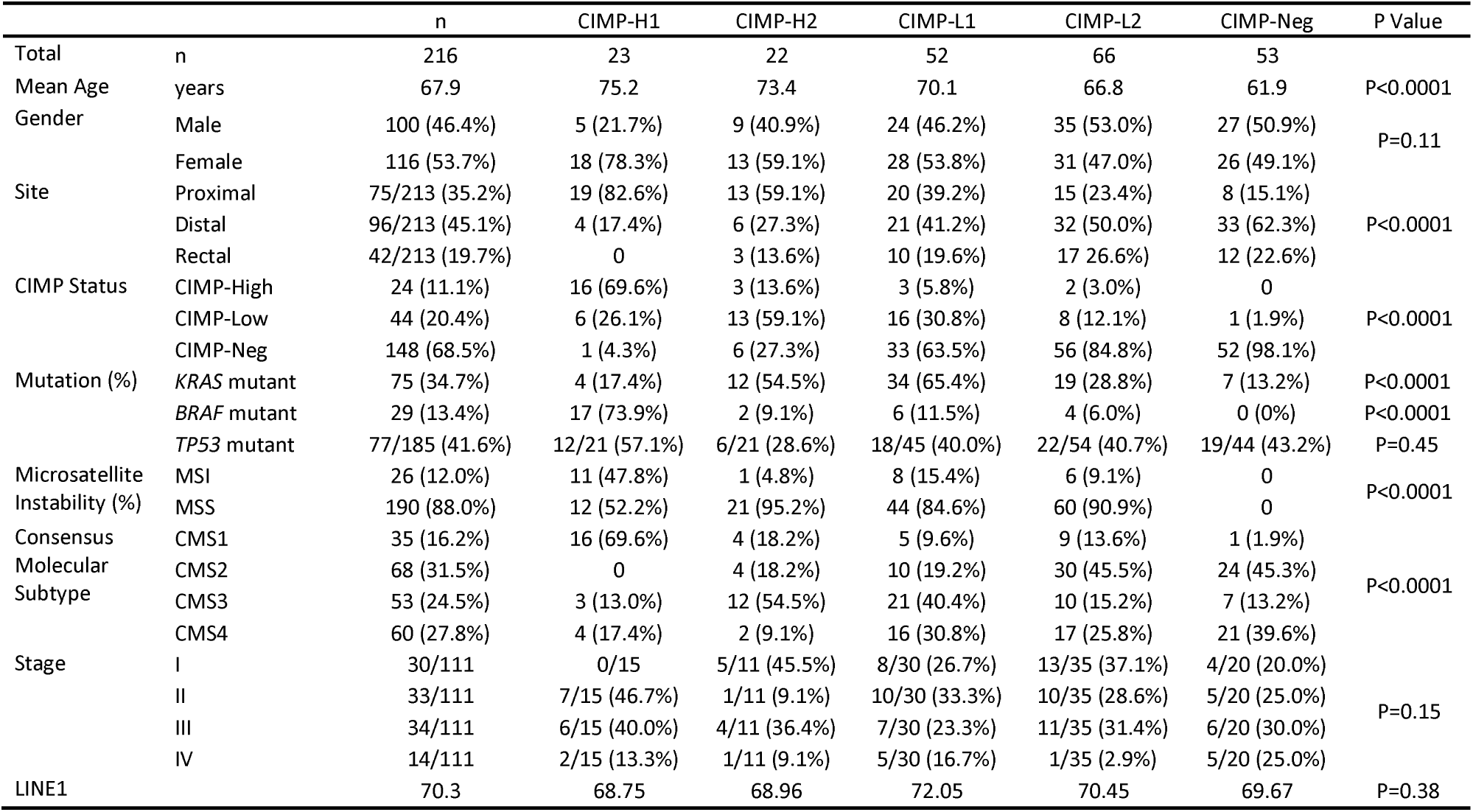
Clinicopathological details of the 216 colorectal adenocarcinomas as stratified for methylation based CIMP clustering, measured on Illumina HM450 arrays, using the 5,000 most variable CpG sites that were not hypermethylated in normal mucosal tissue.

### Methylation-based clustering reveals five subtypes of colorectal cancer with distinct clinical and molecular features

We examined the extent and spectrum of DNA methylation changes in these 216 colorectal cancers using Illumina HumanMethylation450 BeadChip arrays. Five clusters were identified by RPMM clustering (Figure 1). These included two clusters with high levels of methylation that we have designated as CIMP-H1 and CIMP-H2; two clusters with intermediate levels of methylation, CIMP-L1 and CIMP-L2; and a single cluster with low levels of methylation, CIMP-Neg. There was a significant stepwise increase in age between clusters concordant with increasing genomic methylation (CIMP-Neg: 61.9 years, CIMP-L2: 66.8 years, CIMP-L1: 70.1 years, CIMP-H2: 73.4 years, CIMP-H1: 75.2 years, P<0.0001) (Table 1).

**Figure 1:**
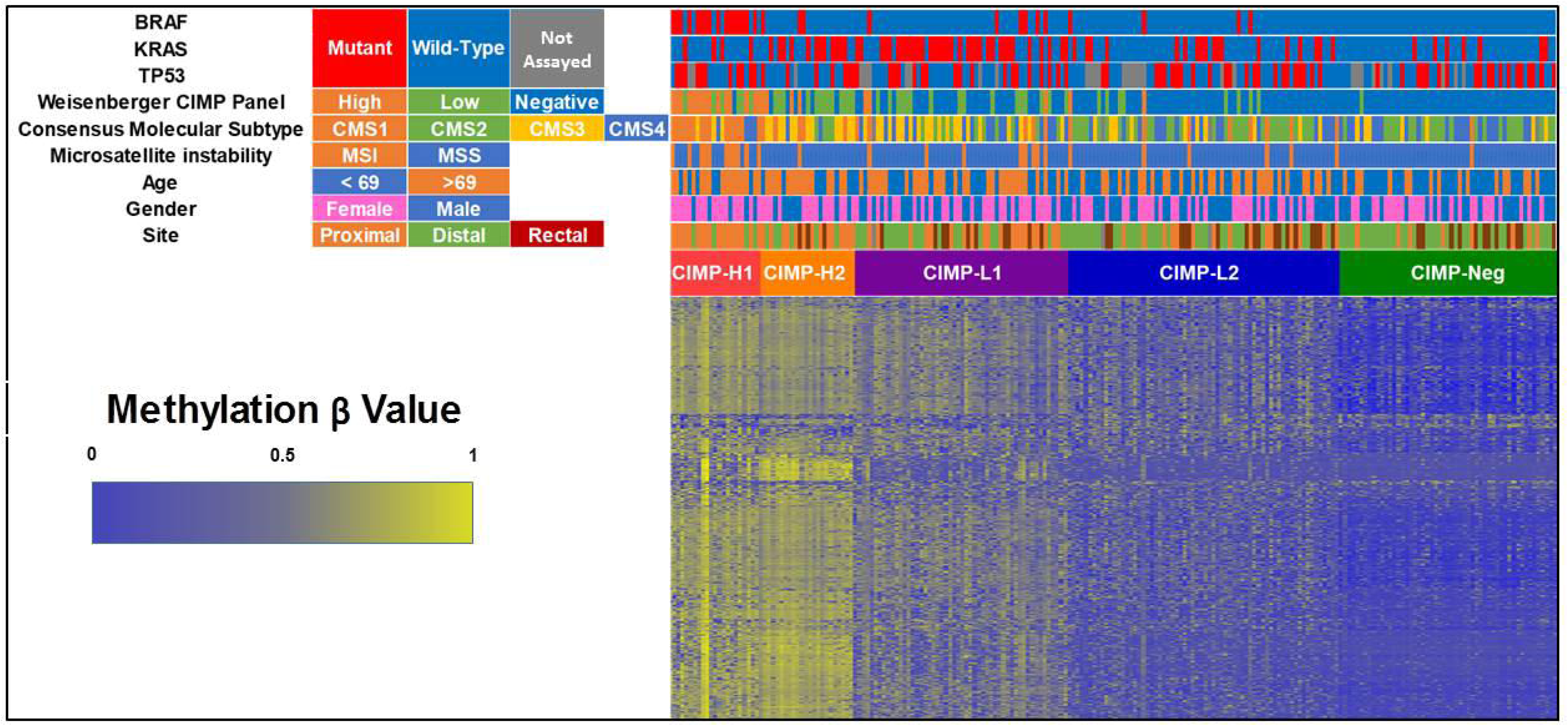
Methylation heatmap of unselected 216 colorectal cancer using the 5,000 most variable beta values in CpG sites that were not hypermethylated in normal mucosal tissue. Clustering was performed using the RPMM R package. Clustering revealed 5 distinct clusters, termed CIMP-Hl, CIMP-H2, CIMP-Ll, CIMP-L2, and CIMP-Neg.

The CIMP-Hl subgroup comprised 23/216 (10.6%) of all cancers and was enriched for female patients (18/23, 78.3%, P<0.0001) and tumours located proximal to the splenic flexure (19/23, 82.6%, P<0.0001). There were no rectal cancers were in the CIMP-H1 subgroup. We observed no differences in stage of cancer at diagnosis and methylation cluster. The CIMP-H1 cluster was strikingly enriched for cancers with features characteristic of serrated neoplasia, including *BRAF* mutation (17/23, 73.9%, P<0.0001), CIMP-H status determined using the Wiesenberger marker panel (16/23, 69.6%, P<0.0001), MSI (11/23, 47.8%, P<0.0001) and consensus molecular subtype CMS1 (16/23, 69.6%, P<0.0001) (Table 1, Figure 1). *TP53* was mutated in 12/21 (57.1%) CIMP-H1 cluster cancers. Of these, 4 were MSI and 8 were microsatellite stable.

CIMP-H2 cluster cancers also frequently arose in the proximal colon (13/22, 59.1%) of females (13/22, 59.1%). These cancers were predominantly microsatellite stable (21/22 (95.2%), *KRAS* mutant (12/22 (54.5%), CIMP-Low as determined by the Weisenberger panel (13/22, 59.1%) and consensus molecular subtype CMS3 (12/22, 54.5%). The CIMP-L1 cluster was also enriched for *KRAS* mutant cancers (34/52, 65.4%, P<0.0001) and had an over-representation of CMS3 (21/52, 40.4%) and CMS4 (16/52, 38.8%) molecular subtypes. The CIMP-L2 and CIMP-negative clusters were predominantly distal or rectal and most likely to CMS2 or CMS4 (Table 1, Figure 1).

### The colorectal cancer methylome is altered in comparison to normal mucosa

We identified differentially methylated probes in each cluster compared to 32 normal mucosal samples that matched a subset of cancers in the unselected series (Table 2, Supplementary Data 1). In all 4 CIMP clusters (CIMP-H1,-H2,-L1 and-L2), the number of differentially hypermethylated CpG sites greatly exceeded those that were *hypo*methylated (Table 2). By contrast, in the single CIMP-negative cluster, hypomethylation was more common than hypermethylation. Probe hypermethylation was most frequent in the CIMP-Hl cluster, including 21,168 hypermethylated probes occurring within 5,165 unique CpG islands. Of these, 4333 were also hypermethylated in CIMP-H2, whilst 832 were uniquely hypermethylated in CIMP-H1. An additional 523 CpG islands were uniquely hypermethlated in the CIMP-H2 cluster relative to CIMP-H1. The highest number of hypomethylation events was seen in the CIMP-H2 cluster compared to all other clusters (P<0.0001), with the majority occurring in open sea regions of the genome.

**Table 2:** Distribution of differentially hypermethylated probes in reference to CpG Islands versus normal mucosal tissue. Cancers are stratified for CIMP Clustering. Differential methylation was deemed as an absolute beta value change of >0.2 and an FDR corrected P Value <0.01 compared to 32 Normal. The ‘+’ symbol refers to differential hypermethylation. The symbol referring to differential hypomethylation.

| CpG Location | CIMP-H1 |  | CIMP-H2 |  | CIMP-L1 |  | CIMP-L2 |  | CIMP-Neg |  |
| --- | --- | --- | --- | --- | --- | --- | --- | --- | --- | --- |
|  | + | - | + | - | + | - | + | - | + | - |
| Island | 21168 | 204 | 19832 | 426 | 11420 | 118 | 5756 | 127 | 760 | 162 |
| South Shore | 3240 | 586 | 3066 | 1359 | 1280 | 426 | 523 | 284 | 78 | 242 |
| North Shore | 4808 | 890 | 4729 | 1885 | 2129 | 617 | 928 | 420 | 187 | 346 |
| South Shelf | 235 | 743 | 189 | 1620 | 85 | 574 | 50 | 331 | 19 | 238 |
| North Shelf | 286 | 738 | 269 | 1660 | 98 | 591 | 62 | 342 | 37 | 246 |
| Sea | 2098 | 8396 | 1810 | 15575 | 664 | 6812 | 307 | 4189 | 109 | 3428 |
| Total | 31835 | 11557 | 29895 | 22525 | 15676 | 9138 | 7626 | 5693 | 1190 | 4662 |

### CIMP-Hl and CIMP-H2 cancers can be delineated by expression profiles

This is the first study sufficiently powered to segregate CIMP-High cancers into two clinically and molecularly distinct subgroups. To examine the extent to which CIMP-H1 and CIMP-H2 are transcriptionally distinct, we analysed differential expression for each cluster with respect to normal mucosa using Illumina HT-12 Expression arrays (Supplementary Table 1). We then performed single sample gene set enrichment analysis (16) to evaluate enrichments in the Hallmark gene set (17) in individual samples (FDR corrected P<0.05). We identified 10 gene sets significantly enriched in CIMP-H1 cancers, 7 of which were related to the immune response (Figure 2). The bile acid metabolism gene set was significantly enriched in CIMP-H2 cancers.

**Figure 2:**
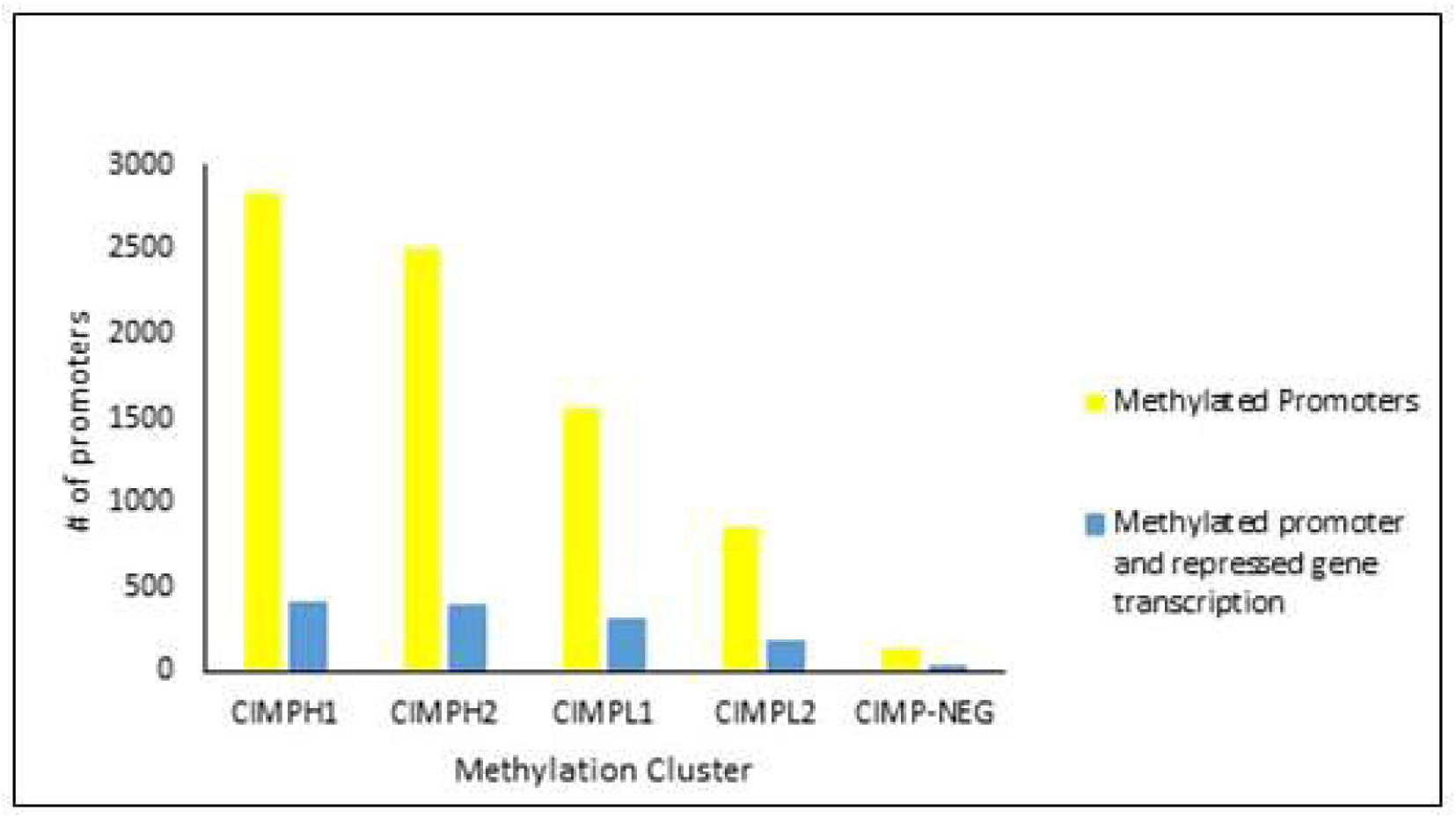
Number of differentially methylated Promoters in each CIMP cluster versus the cohort of normal mucosal samples. The number of Promoters that were methylated, and had a corresponding decrease in transcription is shown, for each cluster, in blue.

### Relationship between promoter hypermethylation and gene transcriptional activity

To determine the frequency to which DNA hypermethylation in promoter regions Controls transcription of downstream genes, we examined the transcript levels for genes where the promoter was hypermmethylated relative normal mucosa (Supplementary Table 2). Although promoter methylation was most common in CIMP-H1 and CIMP-H2 clusters (Figure 3A), these subgroups had the lowest proportion of genes where hypermethylation correlated with reduced transcript expression (14.2% and 15.8%, respectively). This inverse relationship continued for CIMP-L1 (19.2%), CIMP-L2 (20.6%) and with the CIMP-negative cancers having reduced transcription in 22.7% of hypermethylated Promoters (P <0.0001, Figure 3B).

**Figure 3:**
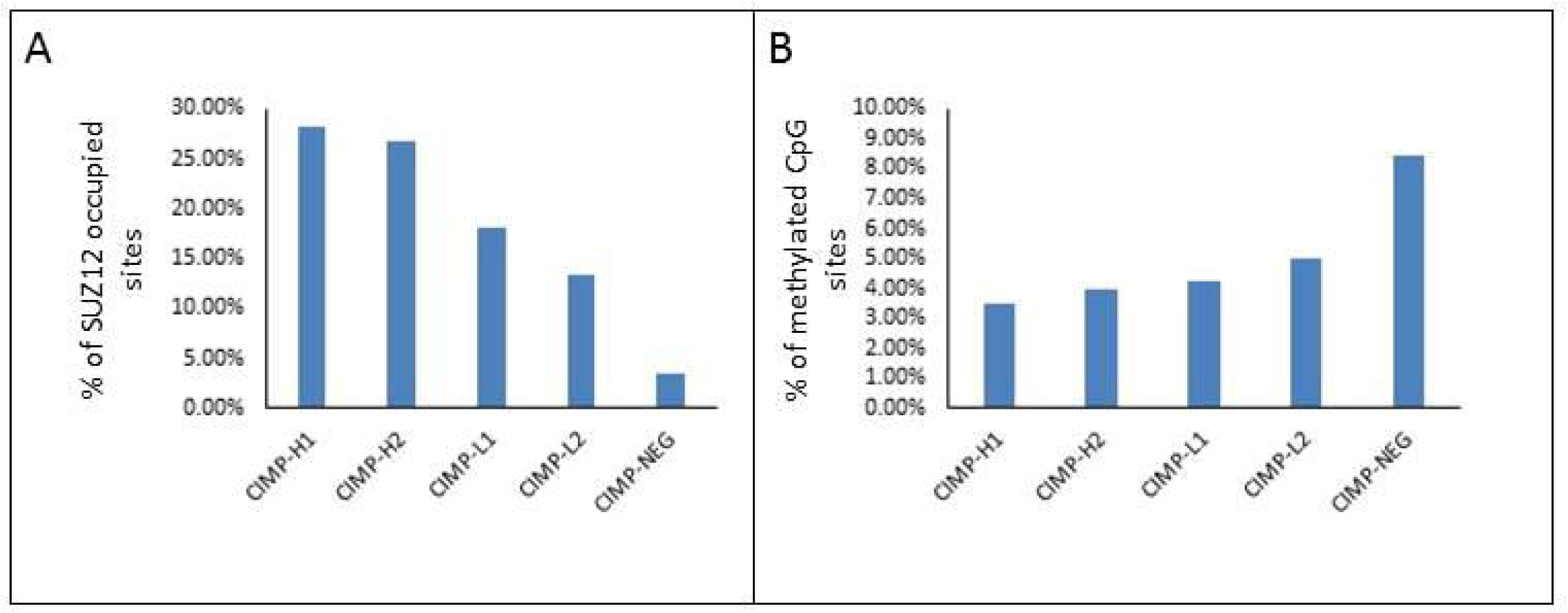
A: Proportion of SUZ12 occupied regions in hESCl cells that contained hypermethylated probes in respective CIMP Clusters. B: Proportion of differential hypermethylation events that overlapped with SUZ12 occupied regions.

### Polycomb-Repressive Complex 2 occupancy at hypermethylated CpGs is inversely correlated with global hypermethylation

SUZ12 occupancy is a Surrogate for polycomb-repressor complex 2 occupancy and in embryonic stem cells this has been shown to associate with transcriptional repression of hypermethylated loci (6, 18). Consistent with this, we observed an increase in SUZ12 occupied sites with increasing CIMP cluster (P<0.0001, Figure 4A). We further observed an inverse association between proportion of hypermethylated loci genes that overlapped with SUZ12 occupied sites with increasing CIMP cluster (PO.OOOl, Figure 4B). This further supports our finding that whilst DNA hypermethylation occurs more frequently with increasing CIMP cluster, these methylation events are more likely to result in gene silencing in CIMP-negative cancers.

**Figure 4:**
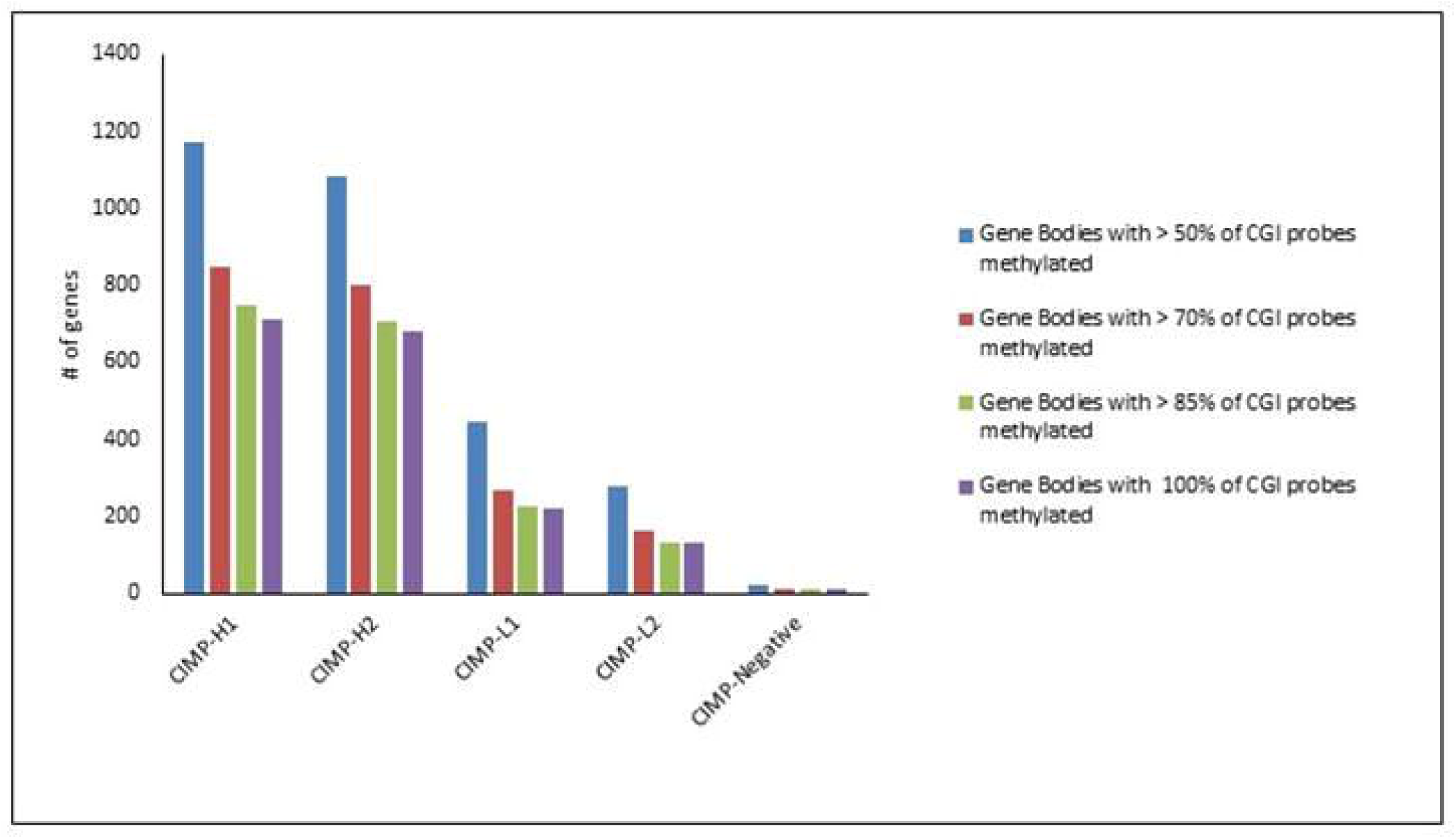
Number of differentially hypermethylated gene bodies stratified for CIMP Cluster and the proportion of CpG island probes within the gene body that are hypermethylated.

### CIMP-Hl and CIMP-H2 promoter methylation is defined by the enrichment of distinct transcription factor binding sites

Transcription factor binding sites often contain CpG sequences and therefore are a target of DNA methylation, which may explain some of the effects of methylation on transcription. To explore whether DNA methylation is targeted to specific transcription factor binding sites we performed an enrichment analysis using the CentriMo (19) tool to examine the 2kb region immediately upstream of hypermethylated genes. There were 128 significantly enriched binding sites that overlapped in CIMP-H1 and CIMP-H2 cancers. An additional 323 sites were uniquely enriched in CIMP-H1 cancers and an additional 330 sites in CIMP-H2 cancers. SMAD4 and FOXP3 (adjusted P Value: 1.2xl0^-24^ and 4.1xl0^-23^, respectively) were the most significantly enriched motifs in CIMP-H1 cancers. SPDEF, FLI1 and NKX6 (adjusted P Value: 7.2xlO^-30^, 1.1×10^-16^, 3.5×10^-16^, respectively) were most significantly enriched in CIMP-H2 cancers. Supplementary Table 3 presents enriched consensus binding sites that were exclusive to CIMP-H1 and CIMP-H2.

**Table 3:** Overlap between genes marked by the PRC2 complex and H3K27Me3 in hEScells and genes which undergo significant gene body methylation in colorectal cancer development. Overlap fraction represents the gene bodies that are methylated (k) divided by the number of genes marked by each respective mark in hEScells (K) (k/K). The FDR corrected P value was obtained through modeling a hypergeometic distribution using the compute overlaps tool on the GSEA web portal using the Benporath gene sets, which were obtained though ChIP-on a Chip analysis of human embryonic stem cells.

| Gene Set Name | CIMP-H1 |  | CIMP-H2 |  | CIMP-L1 |  | CIMP-L2 |  |
| --- | --- | --- | --- | --- | --- | --- | --- | --- |
|  | Overlap Fraction | FDR p-value | Overlap Fraction | FDR p-value | Overlap Fraction | FDR p-value | Overlap Fraction | FDR p-value |
| BENPORATH_ES_WITH_H3K27ME3 | 30.59% | 1.34E-280 | 31.04% | 2.50E-300 | 13.06% | 6.11E-122 | 8.50% | 1.60E-78 |
| BENPORATH_EED_TARGETS | 30.70% | 3.91E-267 | 31.07% | 1.12E-284 | 12.81% | 8.75E-112 | 8.66% | 8.47E-77 |
| BENPORATH_SUZ12_TARGETS | 30.92% | 5.05E-264 | 30.73% | 9.67E-273 | 12.91% | 1.29E-110 | 8.48% | 2.02E-72 |
| BENPORATH_PRC2_TARGETS | 37.27% | 1.04E-218 | 38.04% | 8.59E-235 | 16.41% | 4.56E-98 | 11.04% | 2.05E-66 |

### CpG Island methylation within gene bodies is targeted to cancer related pathways, and occurs more frequently in genes associated with extracellular matrix organization

Gene body methylation is positively correlated with gene expression (20). We examined hypermethylation in gene body CpG islands, defined where >50% of probes in the CpG island were hypermethylated relative to normal (P<0.01) and there was a mean absolute difference in beta values versus normal of >0.2. Gene body CpG island hypermethylation was most prevalent in the CIMP-H1 subgroup and this reduced concordant with reducing global methylation changes (Figure 5). GO pathway enrichment analysis was performed using the Reactome Pathway Gene Sets to determine whether gene body methylation was targeting pathways relevant to carcinogenesis. In CIMP-H1, CIMP-H2 and CIMP-L1 there was a shared underrepresentation of genes regulating the cell cycle (Fold Enrichment: 0.20, 0.25 and 0.001, P=2.9xl0^-7^, 7.4xl0^-6^, 3.2xl0^-2^, respectively). There was an overrepresentation of genes involved in extracellular matrix organization between CIMP-H1, CIMP-H2, CIMP-L1, and CIMP-L2 (Fold Enrichment: 2.87, 2.89, 3.47 and 4.24, P=4.9xl0^-9^, 1.3xl0^-8^, 3.3xl0^-6^, 4.8xl0^-3^, respectively). Supplementary Table 4 presents the pathways that were significantly over and underrepresented in gene body methylation in each CIMP group.

**Figure 5:**
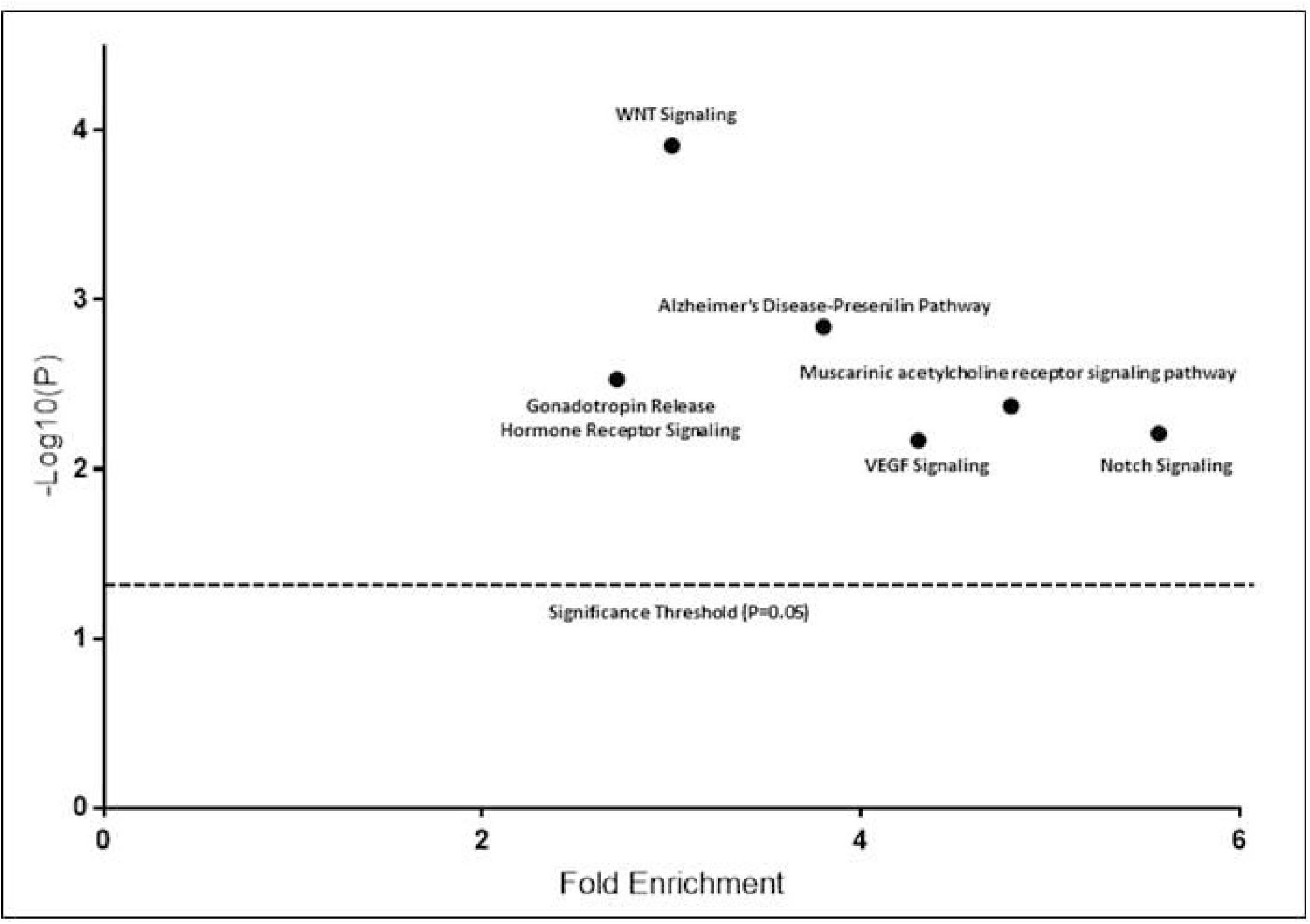
Pathways significantly enriched for genes that contained CpG islands that were devoid of methylation in both CIMP-H clusters.

### Gene bodies of Wnt pathway antagonists are resistant to methylation

We further explored gene bodies that were unmethylated, but had >10 CpG island probes, and performed pathways analysis to identify pathways that were devoid of gene body methylation. There were six pathways that were significantly enriched amongst these genes, including the WNT signaling pathway, the VEGF signaling pathway and the Notch signaling pathway (Figure 6). The WNT signaling pathway was most heavily enriched. *PCDHA6, PCDHGA2, PCDHA7* and *PCDHA2,* which contained 36, 15, 10, and 20 gene body CpG island probes were all unmethylated. These protocadherins have been implicated in regulation of the WNT Signal and may act as tumour suppressor gene. Likewise *AXIN1,* a gene critical to the β-catenin destruction complex, contained 11 unmethylated intragenic CGI probes. *TCF3,* a WNT pathway repressor, contained 19 unmethylated intragenic CGI probes. These data indicate possible role for gene body demethylation in WNT signaling regulation in CIMP-H cancers.

**Figure 6:**
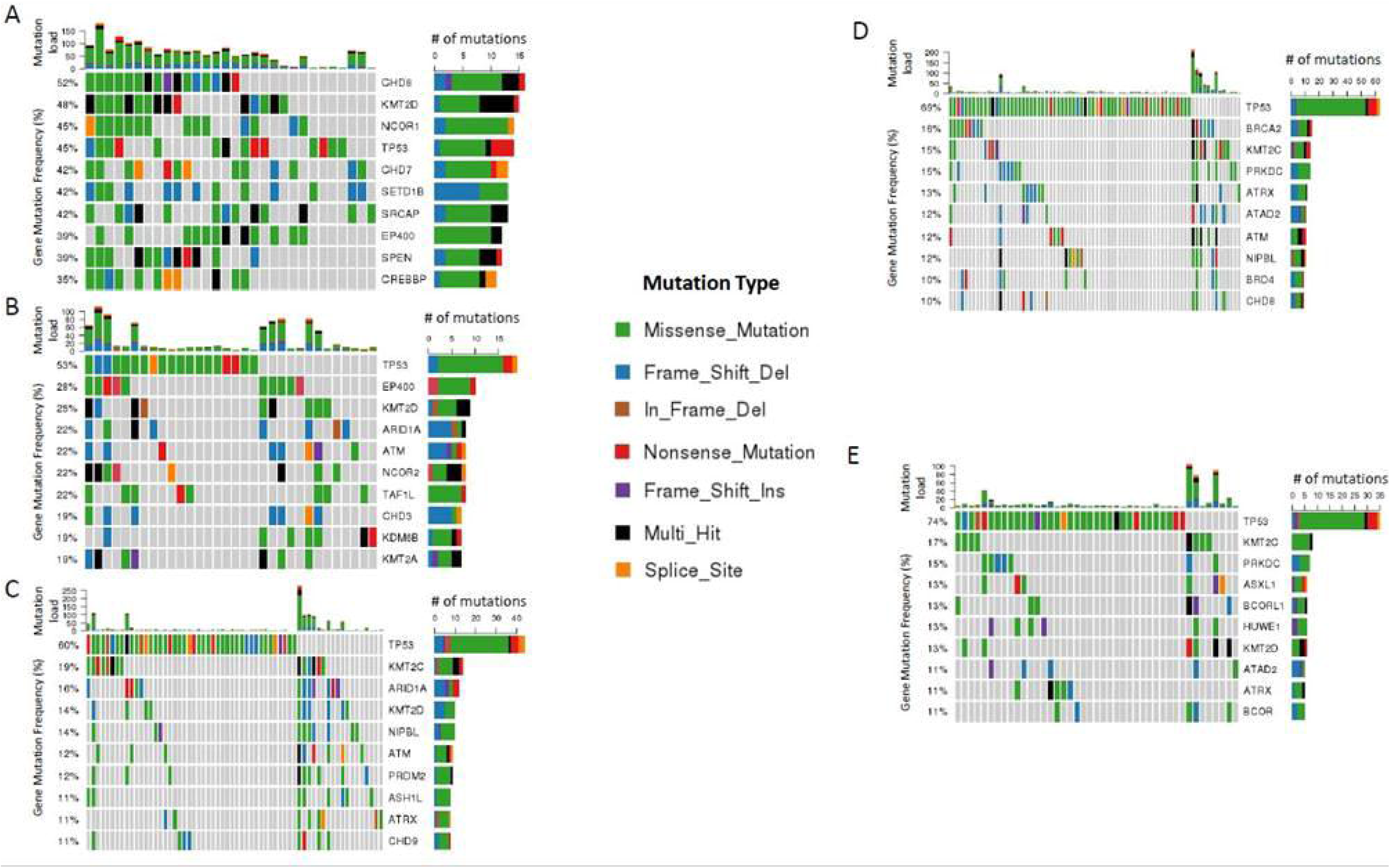
Summary of the most frequently observed epigenetic regulatory gene mutations. Gene mutation frequency for each loci is shown in the Y axis of the central plot. The barplot on the uppermost plot represents the total number of mutations in epigenetic regulator genes within each sample. A) CIMP-Hl B) CIMP-H2 C) CIMP-Ll D) CIMP-L2 E) CIMP-Neg

### Oncogenes arefrequently targeted for gene body hypermethylation

In CIMP-H1 cancers, the gene bodies of 47 annotated oncogenes were significantly hypermethylated, including a subset of 31 oncogenes where all gene body probes were hypermethylated (eg *ERBB4* and *BCL2,* Supplementary Table 5). Whilst 38 oncogenes were methylated in the gene bodies of both CIMP-H1 and CIMP-H2 cancers, there were 9 genes exclusively hypermethylated in gene bodies in CIMP-H1 cancers (*NKX2-1, BCL2, SALL4, PRDM8, KIT, LAPTM4B, MERTK, CYP24A1* and *WNT1)* and 9 exclusively hypermethylated in CIMP-H2 cancers (*WWTR1, RET, PRDM6, PAX8, GRM1, CXCR4, SLC12A5, PPP1R14A* and *BMP7).*

### Loci marked by the PRC2 complex in human embryonic stem cells are prone to gene body methylation during cancer development

PRC2 marking in human embryonic stem cells has previously been shown to overlap significantly with promoter hypermethylation in colorectal cancers (6). We hypothesized that a similar phenomenon would occur with regards to gene body hypermethylation. In CIMP-H1 and CIMP-H2 cancers, 30.59% and 31.04% of loci marked with H3K27me3 in hEScells developed significant gene body hypermethylation (Table, P=1.34xlO^-280^ for CIMP-Hl and P=2.5xlO^-300^ for CIMP-H2 overlap). We observed a lesser, but still highly significant overlap between H3K27me3 marked loci and gene body methylation in CIMP-L1 (13.1%, P=6.11×10^122^) and CIMP-L2 (8.5%, P=1.6xl0^-78^) cancers but did not observe any correlation in CIMP-Neg cancers, which is likely due to the scarcity to which gene body methylation occurs in these cancers. We observed similar overlaps for EED targets, SUZ12 targets and PRC2 targets.

### Epigenetic regulator gene mutations are common in The Cancer Genome Atlas cancers

Mutations in epigenetic modifier genes have previously been shown to modulate transcriptional profiles in cancer (15). We assessed the mutational frequency of 719 epigenetic regulator genes in TCGA Colon Adenocarcinoma cancers (21) that we had assigned to CIMP clusters using a machine learning approach based on the same subset of probes used in our unselected series. To test the specificity of our model we compared known clinical and mutational data for *BRAF* and *KRAS* in each TCGA cohort CIMP clusters with our unselected series. No significant differences were identified, supporting our confidence in the model (Adjusted P value ränge: 0.58-1). We then used overall TCGA survival data to assess the impact of CIMP clusters on prognosis. There was no significant difference in survival between the CIMP clusters.

In the TCGA dataset, all cancers had at least 1 mutation in an epigenetic regulator gene (Supplementary Table 6). Figure 7 shows the most commonly mutated epigenetic regulators in each cluster. Mutations were least common in cancers classified as CIMP-Neg, with increasing global methylation being associated with a concordant increase in epigenetic mutational load (Figure 8, P<0.0001). However, when we examined epigenetic mutation frequency in relation to microsatellite instability, there was no significant relationship between CIMP cluster and epigenetic mutation frequency (One-way ANOVA for CIMP clusters trichotomized for MSI status: P=0.91, P=0.99 and P=0.61 for differences between CIMP clusters in MSI-H, MSI-L and MSS, respectively), indicating that the differences observed between CIMP clusters may be driven by the increasing frequency of microsatellite instability in CIMP clusters with higher genomic methylation. 591 genes were mutated in at least one instance in the CIMP-H1 cluster. CIMP-L1 and CIMP-L2 mutated a wider array of genes (552 and 522, respectively) when compared to CIMP-H2 (452 genes mutated), despite having a lower average number of mutations in epigenetic regulators per sample.

**Figure 7:**
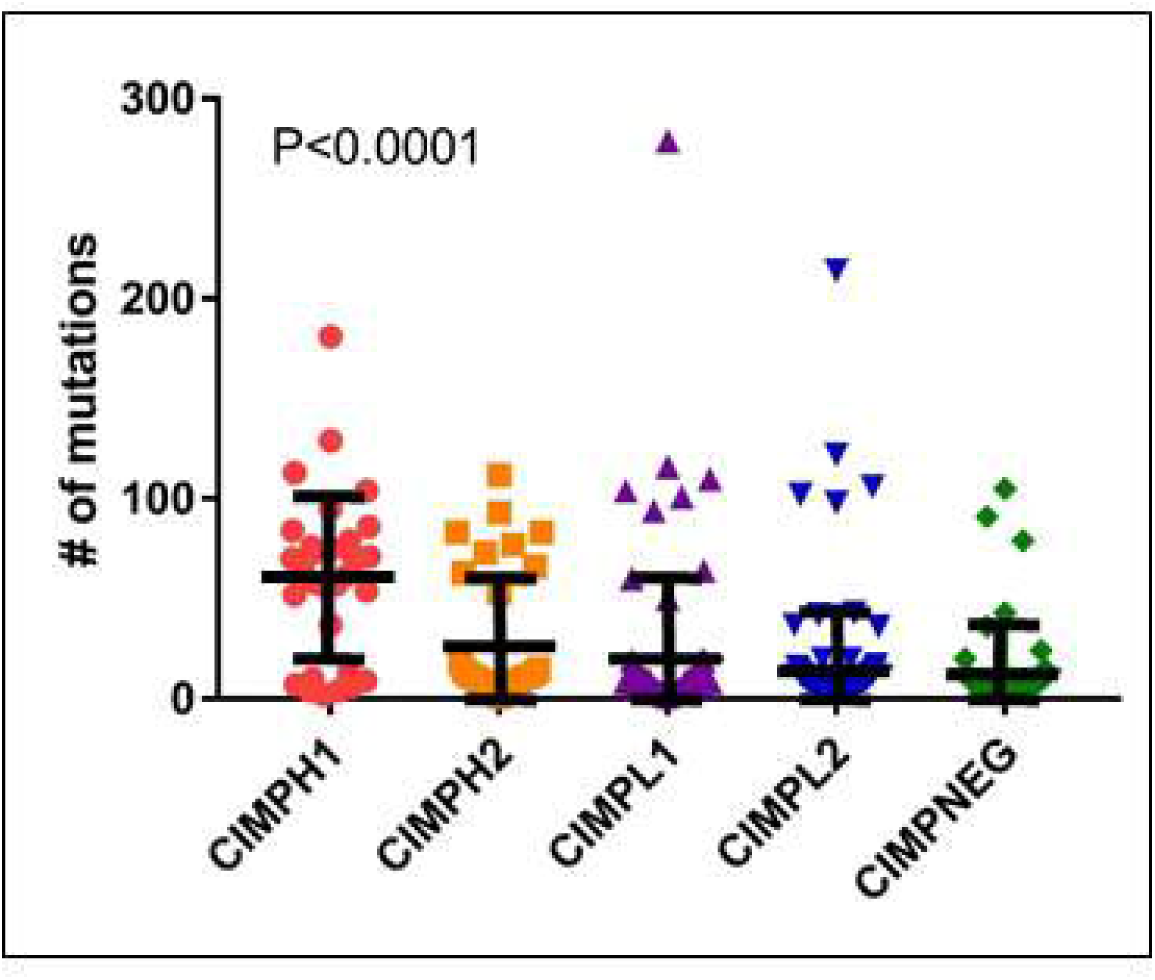
Number of mutations in epigenetic regulator genes as stratified for CIMP cluster. ANOVA was used for statistical analysis.

**Figure 8:**
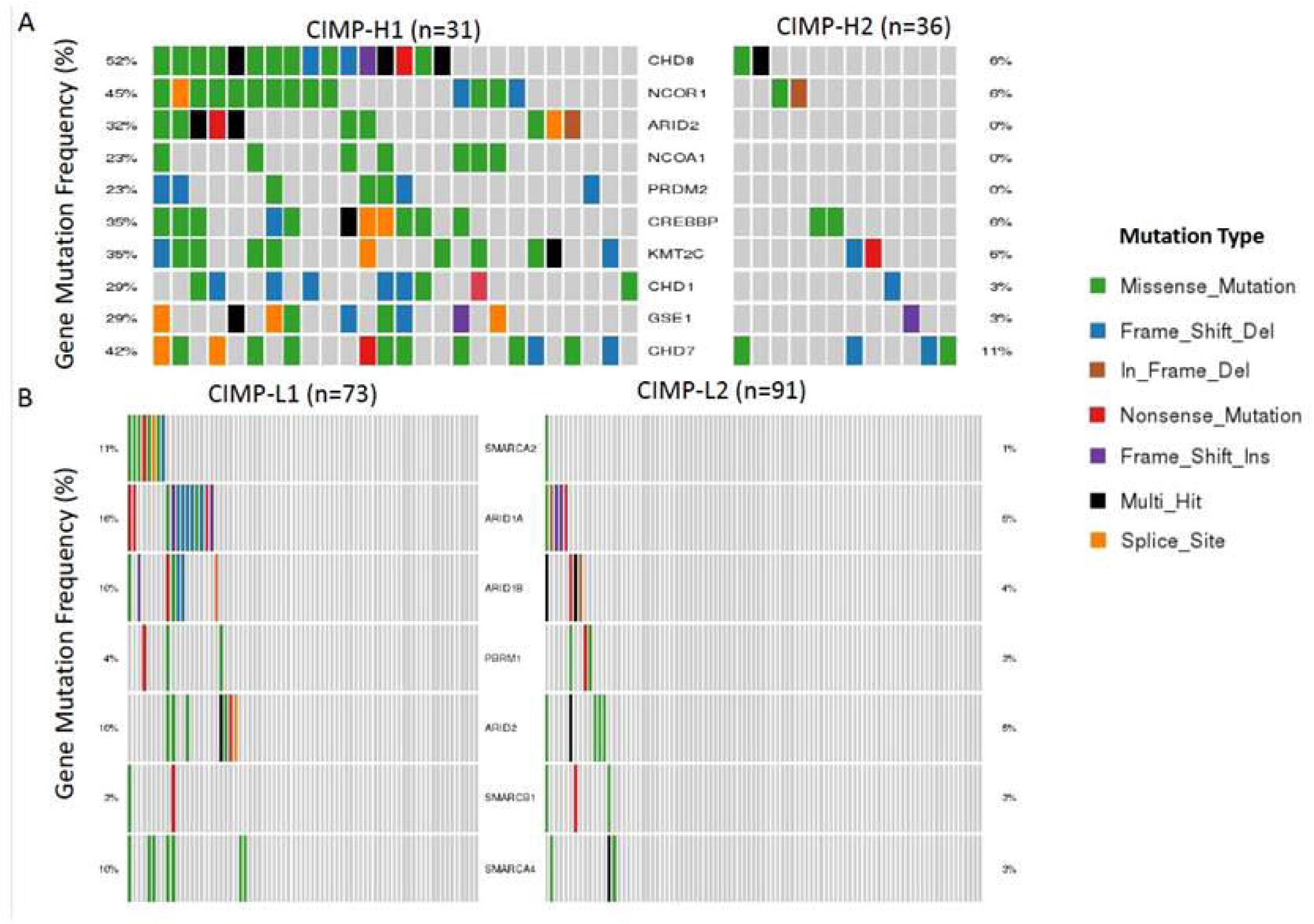
A) Co-Oncoplot of the 10 most significantly differentially mutated genes between CIMP-Hl and CIMP-H2. Differentially mutated genes were determined using 2×2 contingencies and the fishers exact test for each gene. B) Co-Oncoplot comparing mutational frequency of SWI/SNF complex members between CIMP-Ll and CIMP-L2.

### CIMP-Hl and H2 clusters have a similar mutational patterns in epigenetic regulator genes

We sought to elucidate differences in somatic mutational profiles amongst epigenetic regulators between CIMP-H1 and CIMP-H2 cancers. In total, 626 genes were mutated in either CIMPH1 or CIMPH2. 66.6% (417) of these genes were mutated in at least one cancer of each CIMP-H cluster. Only 5.6% (35) were exclusively mutated in CIMP-H2 in comparison to CIMP-H1. By contrast, 27.8% (174) genes were exclusively mutated in CIMP-H1 cancers versus CIMP-H2. 52 genes were mutated significantly more frequently in CIMP-H1 cancers when compared with CIMP-H2 cancers (Figure 9A depicts the top 10 differentially mutated genes). The overall mutational load in CIMP-H1 was higher than in CIMP-H2. As this group was enriched for microsatellite unstable cancers, it is likely that the genetic instability is driving the mutational differences between CIMP-H1 and-H2. Genes significantly more commonly mutated in CIMP-H1 compared to CIMP-H2 include the members of the chromodomain helicase family *CHD8* (OR 17.29, 2.2xl0^-5^), *CHD1* (OR 13.82, P=0.004) and *CHD7*(OR 5.62, P<0.005). Other genes exclusively associated with CIMP-H1 in comparison to CIMP-H2 included *ARID2* (PcO.OOl), *NCOA1* (P=0.003) and *PRDM2* (P=0.003).

**Figure 9:**
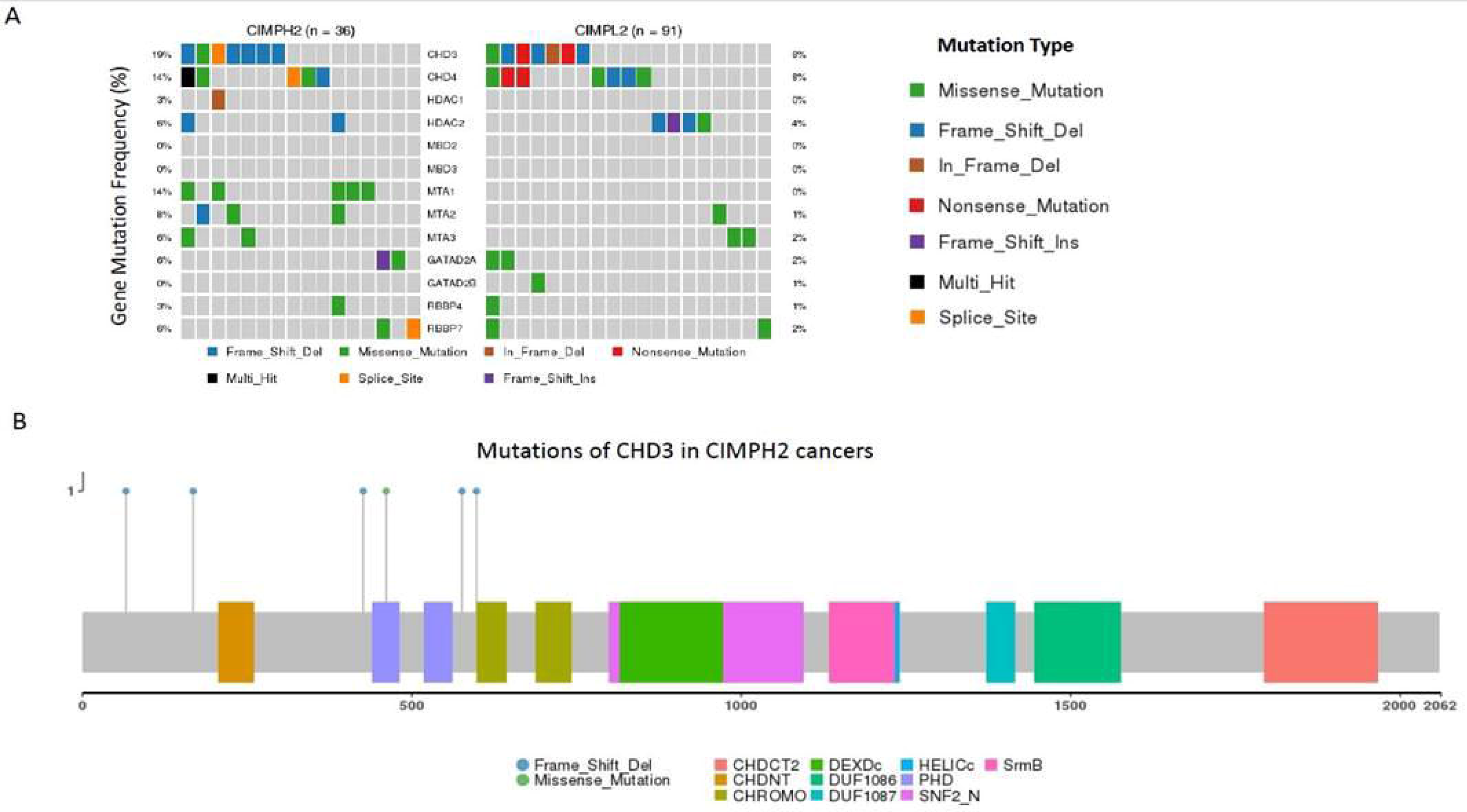
A) Co-Oncoplot comparing the mutational frequency of NuRD complex subunits between CIMP-H2 and CIMP-L2. B) Location of mutations observed in the CHD3 gene in cancers of the CIMP-H2 cluster. The X-Axis indicates the amino acid position.

### Epigenetic regulator gene mutation exclusivity supports the dichotomization of CIMP-L clusters

We examined the frequency and differential mutation rates of epigenetic genes in CIMP-L1 and CIMP-L2 cancers. Mutations in 12 genes were significantly associated with either CIMP-L1 or CIMP-L2. *DNAJC2,* an epigenetic modulator of polycomb-repressed genes, was exclusively mutated in CIMP-L2 cancers and not in CIMP-L1. Four of eight mutations in *DNAJC2* were truncating. *DNAJC2* has been associated with oncogene induced senescence via the *INK4/ARF* cascade (22), and therefore inactivating mutations in this gene may be associated with overcoming oncogene induced senescence. By contrast, we observed significantly greater mutations in subunits of the chromatin remodeling SWI/SNF complex in CIMP-L1 cancers compared with CIMP-L2 (25/73 versus 15/91, p= 0.01) (Figure 9B). The SWI/SNF complex is one of four chromatin remodeling complexes responsible for stimulating gene expression in different cellular contexts. Synthetic lethality in the SWI/SNF complex has previously been established (23) and notably, in CIMP-L1 cancers mutations are frequently truncating. Hence, CIMP-L1 cancers may be more vulnerable to treatments targeting the other element of the SWI/SNF complex. To test whether one SWI/SNF mutation confers dependency on other SWI/SNF subunits *in vitro,* we correlated exome-capture data from 15 cell lines (24) with cell line dependency data from Meyers et al (25). Five cell lines had an *ARID1A* truncating mutation and these were significantly more dependent on *ARID1B* expression for survival (0.31 vs 0.06, P<0.05).

### NuRD complex genes are frequently disrupted in CIMP-H2 and CIMP-L2 cancers

CIMP-H2 and CIMP-L2 cancers frequently harbored truncating mutations in *CHD3/4,* which encode critical subunits of the nucleosome remodeling and deacetylase (NuRD) complex. The NuRD complex is unique in that it is the only chromatin remodeling complex capable of deacetylating histones (26). High-impact mutations were present in 11/36 (30.5%) of CIMP-H2 cancers and 11/91 (12%) of CIMP-L2 cancers (P=0.01, Figure 10A). Critically, truncating *CHD3* mutations in CIMP-H2 occurred before important functional domains, with 4 high impact mutations occurring before the PHD-Finger domain, which is important for recognition of the lysine methylation histone H3. A further 2 mutations occurred prior to the sequence encoding the chromodomain (Figure 10B). Intriguingly, we only observed 1 truncating mutation in each of *CHD3* and *CHD4* in CIMP-H1, despite the relatively higher instances of truncating mutation in cancers in other CIMP groups.

## Discussion

Remodeling of the epigenome is fundamental to colon cancer progression and is a key driver of serrated pathway cancers that typically display the CpG Island Methylator Phenotype. We aimed to better understand the extent of this phenotype, the spectrum of DNA methylation sites targeted and the consequences on gene expression. Through interrogation of the largest unselected series of colorectal cancers to date, using genome-scale technology, we identified five clinically and molecularly distinct DNA methylation clusters. We also identified a striking increase in patient age with increasing DNA methylation cluster, highlighting the importance of the aging colon in the development of serrated colorectal neoplasia.

Hinoue and colleagues previously reported the presence of four colorectal cancer methylation subgroups by assessing 125 colorectal cancers using Illumina 27K DNA methylation arrays (27). In the present study, we have considerably increased the power to assess subgroups based on differential methylation by studying 216 unselected cancers using the Illumina 450K DNA methylation platform. A major difference in our findings is the Segregation of CIMP-H cancers (approximately 22% of the Hinoue *et al* cohort) into two subgroups. Together, our CIMP-H1/H2 clusters represent 21% of our unselected cohort. The dichotomization of these CIMP-H cancers identified a homogeneous subgroup of 23 CIMP-Hl cancers with an average age of 75 years, striking over-representation of female gender and *BRAF* mutant cancers arising in the proximal colon. There were no rectal cancers in the CIMP-H1 group compared to 35% of the CIMP-H2 group. CIMP-H2 cancers preferentially activated the MAPK pathway by mutation of the *KRAS* oncogene. Together, these data suggest that CIMP-H1 cancers are more likely to arise from sessile serrated adenomas whilst CIMP-H2 cancers arise from traditional serrated adenomas (28).

We observed a consistent increase in patient age with CIMP cluster, from 62 years in CIMP-Neg cancers to 75 years in CIMP-H1 cancers. This is in contrast to the Hinuoe study (6). The variance in our assay was mostly contained in uniquely mapping probes that were not present in the 27K array employed by Hinuoe et al. Numerous studies have demonstrated age-related methylation in different tissues (9, 29, 30) and we have previously identified hypermethylated loci in the colons of patients even with no history of colonic disease (9). In the present study, we detected a significant correlation between methylation and biological age of the participant. After removal of all probes that were significantly hypermethylated in normal mucosal tissue, we still observed distinct, age linked clustering. It is possible that serrated pathway cancers require age-related methylation ‘seeds’ that spread over time to silence key genes for tumour progression, and that this can be accelerated by activation of the MAPK pathway by oncogenic mutation of *BRAF* or *KRAS.* This hypothesis is supported by our recent finding that *BRAF* mutant sessile serrated adenomas of the colorectum rarely exhibit the classic methylator phenotype until after 50 years of age (31). This is also consistent with our animal model for serrated neoplasia where we observe a slow accumulation of DNA methylation changes over time, however these are dramatically accelerated by mutating *BRAF,* congruent with development of serrated neoplasia (32). This may explain why *BRAF* mutant sessile serrated adenomas are often identified in younger patients, despite the cancers arising from them occurring primarily in older patients (12, 33, 34). Understanding the role of age-related methylation might inform surveillance for younger patients with sessile serrated adenomas.

The striking over-representation of female gender in the CIMP-H1 cluster may relate to hormonal changes that increase the rate of age-related epigenetic drift, the process whereby DNA accumulates methylation over a protracted period due to a reduction in the fidelity of DNMT enzymes. Consistent with this hypothesis, Levine *et al* have shown that menopause increases epigenetic drift (35), and Noreen *et al* demonstrated that hormone replacement therapy reduces epigenetic drift (36).

Differential CpG island and shore hypermethylation were the most frequently observed methylation events in the study. Probes on the north and south CpG shelves, as well as those in the open seas were frequently hypomethylated across most cancers. The implications of hypomethylated CpG dinucleotides outside of CpG islands are unclear. We did not observe any relationship between hypomethylation and gene transcription, however it is possible that hypomethylation of specific regions of the genome may affect chromatin accessibility elsewhere and hence may modulate transcription in a trans-acting manner. Open sea hypomethylation was also the most frequent methylation event in CIMP-Neg cancers. These are predominately conventional pathway cancers with a high degree of chromosomal instability. It is possible that hypomethylation outside of CpG islands may predispose to copy number changes in these cancers (37, 38). Additional studies are necessary to explore the functional implications of shelf and open sea hypomethylation.

There were marked differences in transcriptional deregulation of key cancer-related pathways between methylation clusters. CIMP-H1 cancers activated several immune pathways, including those involved in the interferon response, inflammatory response and complement signaling, consistent with the over-representation of CMS1 cancers in this group. This is likely due to the higher mutational burden in these cancers, largely driven by the increased incidence of epigenetically induced microsatellite instability. CIMP-H2 cancers were uniquely enriched for altered bile acid metabolism, consistent with the previously described relationship between silencing of the farnesoid X bile acid receptor in *KRAS* mutant cancers (39). Bile acids are more concentrated in the proximal colon and metabolism is influence by the gut microbiome (40), which may provide insight into causation of this particular cancer subgroup. Better understanding the role of bile acid signalling in *KRAS* mutant cancers of the proximal colon may have therapeutic implications for this cancer subgroup.

Paradoxically, despite observing less differential methylation, we observed an increase in gene silencing that correlated with promoter hypermethylation in the least methylated cancer clusters. This may indicate that promoter hypermethylation in CIMP-L½and CIMP-Neg cancers is more specifically selected based on a functional advantage in these cancers. Alternatively, the increased frequency of mutations in epigenetic regulators of CIMP-H½ cancers may result in a reduced capacity to induce gene repression at certain loci. This may be due to the loss of a repressive histone modifying enzyme, or mutation of locus specific repressive transcription factors. Methylation alone may be insufficient to induce gene repression in certain instances. Instead, relevant chromatin remodelling and histone modifications, such as the addition of the repressive PRC2 mark, may be required in tandem with methylation changes to reduce gene expression. Indeed, we showed that PRC2 occupancy was most frequently related to transcriptionally repressed and methylated genes in the CIMP-Neg subgroup. We also observed instances of promoter methylation that correlated with increased gene transcription. It is possible that some transcription factors preferentially bind methylated DNA (41), and that binding sites for these transcription factors become available following promoter methylation. These data highlight the importance of the genomic and epigenomic context in which methylation occurs.

A major novel finding of the current study is the discovery that gene body methylation may be a major driver of serrated tumorigenesis, and that this may be mediated by H3K27me3 histone marks. Gene body hypermethylation has recently been correlated with increased oncogene expression (20). Here we identified many well characterised oncogenes, such as *ERBB4* and *BCL2,* with methylation of their gene bodies in CIMP-H½ cancers. We also identified Wnt pathway antagonists that are resistant to gene body methylation, which may limit expression of these tumour suppressor genes. The role of gene body methylation in serrated neoplasia, particularly in the context of epigenetic therapy, requires further investigation.

The epigenome is regulated by proteins that interact with histones or DNA. We assessed the coding sequence of 719 epigenetic regulator genes in the TCGA dataset. The chromodomain-helicase-DNA (CHD) binding protein family was a frequent mutational target in CIMP-H1 cancers. Recently, Fang et al. showed that CHD8 operates in a transcriptional repression complex to direct methylation in the setting of *BRAF* mutation (42). In the current study we showed *BRAF* and *CHD8* mutations were associated with CIMP-H1. Thus these data suggests that *CHD8* mutation may enhance repression complex activity in the setting of *BRAF* mutation, resulting in hypermethylation. Moreover, CHD8 has been associated with the CTCF protein, which is essential for promoter-enhancer looping and regional insulation. *CHD8* mutations may influence CIMP by decreasing the ability of CTCF to insulate regions of the genome, and could encourage methylation spreading throughout the genome (43).

Chromatin remodeling is an essential process whereby Condensed euchromatin is modified in a context-specific manner to give rise to regions of heterochromatin that can be actively transcribed. Chromatin remodelling is driven by a series of complexes that are able to enzymatically catalyze reactions that modify histone tails and, in turn, modulate the accessibility of the chromatin. In mammalian cells five key chromatin modifying complexes predominate. The chromodomain helicase DNA-binding complex (CHD), the IN080 complex, the SWI/SNF complex, ISWI complex and the NuRD complex (44). We examined the frequency of mutations within the coding regions of genes that encode subunits of these complexes. In CIMP-H2 and CIMP-L2 cancers we observed frequent mutations in members of the NuRD complex, which has both chromatin remodeling and histone deacetylation capabilities. Truncating mutations in *CHD3/4* may indicate a therapeutic vulnerability to DNMT inhibitors, as a result of the synthetic lethality of the NuRD complex and the DNMT proteins (45) in CIMP-L1 cancers, despite not having a classical hypermethylator phenotype. Further study is necessary to explore the role of DNMT inhibition in cancers with NuRD complex mutations. Similarly, CIMP-L1 cancers had frequent SWI/SNF subunit mutations. Frameshift mutations in *ARID1A/B* were the most common mutations. It is well established that SWI/SNF mutations confer synthetic lethality upon other subunits. To test this hypothesis we used public colorectal cancer cell line dependency data in conjunction with mutational data, and identified a strong dependency conferred upon ARID1B following genetic perturbation of ARID1A. These data Support the investigation of SWI/SNF inhibitors to exploit synthetic lethality presented by SWI/SNF mutations in CIMP-L1 cancers.

## Conclusion

The past decade has heralded an era where the importance of the cancer epigenome is increasingly recognized, where treatments targeting different epigenetic modifications are entering the clinic and improving patient outcomes. It has become apparent that a comprehensive understanding of the epigenetic drivers of cancer will be crucial in the rational design of clinical trials and the development of precision medicine strategies. Here we have identified five clinically and molecularly distinct subgroups based on a comprehensive assessment of a large, unselected series of colorectal cancer methylomes. In contrast to earlier studies, we identify two CIMP-H clusters which are demarcated by *BRAF* and *KRAS* mutation status. We observe a striking association between genomic methylation and age, which further supports the investigation of the epigenetic clock in serrated neoplasia risk. We identify a novel role for gene body methylation in serrated neoplasia, which may be mediated by H3K27me3 histone marks. Our interrogation of the coding regions of epigenetic regulatory genes shows that they are frequently mutated in colorectal cancers and this is partially influenced by the degree of genomic methylation. Our analyses have identified potentially druggable vulnerabilities in cancers of different methylation subtypes. Inhibitors targeting synthetic lethalities, such as DNMT inhibition for cancers with NuRD complex mutations and SWI/SNF component inhibitors for those with *ARID* mutations, should be evaluated as these agents may be clinically beneficial to certain patient subsets.

## Methods

### Patient samples

Colorectal cancer (N = 216) and matched normal (N = 32) samples were obtained from patients undergoing surgery at the Royal Brisbane and Women’s Hospital, Brisbane, Australia, in a consecutive manner between 2009 and 2012. Tissue was snap-frozen in liquid nitrogen to preserve sample integrity. Written informed consent was obtained from each patient. The study protocol was approved by the Royal Brisbane and Women’s Hospital and QIMR Berghofer Medical Research Institute Research Ethics Committees. The Cancer Genome Atlas (TCGA) colon adenocarcinoma (COAD) exome and methylation data (N = 278) were used for independent validation (21).

### DNA and mRNA extractions

DNA and mRNA were simultaneously extracted from approximately 30 mg of homogenized tissue using the AllPrep DNA/RNA Kit (Qiagen, Australia) in accordance with the manufacturer’s protocols. Double stranded DNA concentration was assessed using the PicoGreen quantitation assay (Molecular Probes, USA). mRNA quality was measured using the Bioanalyzer 2100 platform (Agilent, USA). Microarray analysis was performed on samples with a RNA integrity number of >7.

### Molecular characterization of cancer samples

Cancer sample DNA was analyzed for the *BRAF* V600E mutation using allelic discrimination as previously reported (46). In addition, we assayed mutations in *KRAS* codons 12 and 13, and *TP53* exons 4 to 8 using previously reported methods (47, 48). We assessed CIMP status by methylation-specific PCR using the five-marker panel (CACNA1G, IGF2, NEUROG1, RUNX1 and SOCS1) proposed by Weisenberger et al. (3). Samples were considered CIMP-high if ≥ 3 markers were methylated, CIMP-low if 1 or 2 markers were methylated, and CIMP-negative if no markers were methylated. MSI was assessed using the criteria of Nagasaka et al. (49) where instability in ≥ 1 mononucleotide marker, and ≥ 1 additional, non-mononucleotide marker, using the marker set reported in Boland et al., (50) was indicative of MSI, the remainder being microsatellite stable (MSS). *LINE1* methylation was assessed using pyrosequencing as per Irahara et al. (51). CIMP-high cancers that were both *KRAS* and *BRAF* wild-type at hotspot codons were Sänger sequenced for *BRAF* exons 11 and 15 (exon 11, forward 5’-TTCCTGTATCCCTCTCAGGCA-3’, reverse 5’-AAAGGGGAATTCCTCCAGGTT-3’; exon 15, forward 5’-GGAAAGCATCTCACCTCATCCT-3’, reverse 5’-TAGAAAGTCATTGAAGGTCTCAACT-3’), *KRAS* codon 61 (forward 5’-TCCAGACTGTGTTTCTCCCTTC-3’, reverse 5’-TGAGATGGTGTCACTTTAACAGT-3’), and *EGFR* exon 18 (forward 5’-ATGTCTGGCACTGCTTTCCA-3’, reverse 5’-ATTGACCTTGCCATGGGGTG-3’).

### DNA methylation microarray

Genome-scale DNA methylation was measured using the HumanMethylation450 BeadChip array (Illumina, USA). The BeadChip array interrogates cytosine methylation at >480,000 CpG sites. 500ng of DNA was bisulphite converted using the EZ-96 DNA Methylation Kit (Zymo Research, USA) as per the manufacturer’s protocol. Whole-genome amplification and enzymatic fragmentation was performed on post-treatment DNA, which was subsequently hybridized to the array at 48°C for 16 hours. Arrays were scanned using the iScan System (Illumina, USA).

### Gene expression microarray

Gene expression levels for over 47,000 transcripts were measured for all samples using the HumanHT-12 v3 Expression BeadChip array (Illumina, USA). Total mRNA (500 ng) was reverse-transcribed, amplified and biotinylated using the TotalPrep-96 RNA Amplification Kit (Illumina, USA). The labelled cRNA (750 ng) was hybridized to the array followed by washing, blocking, and staining with streptavidin-Cy3. Arrays were scanned on the iScan System and the data was extracted using GenomeStudio Software (Illumina, USA).

### Data analysis

Methylation microarray data were checked for quality against parameters provided by Illumina using the GenomeStudio Software package. IDAT files were read into the R environment using Limma (52). We used subset-within-array normalization (SWAN) to correct for biases resulting from type 1 and type 2 probes on the array. We filtered probes that had a detection P > 0.05 in > 50% of samples, as well as probes that were on the X or Y chromosome, where the CpG site was within lObp of a single nucleotide polymorphism, or where a probe mapped to the genome ambiguously. At the conclusion of filtering 377,612 probes remained and were used for subsequent analyses.

The recursively partitioned mixed model (RPMM) clustering method (53) was used for unsupervised clustering. In order to capture cancer specific methylation we followed methods employed by based The Cancer Genome Atlas (54). DNA methylation drift with age has been charactarised in a number of different normal and cancerous tissues (10). To limit confounding from methylation that occurs through age probes with a mean β value of >0.3 in normal samples were excluded from clustering analysis 144,542 probes were unmethylated (mean β value <0.3) in normal mucosa, of these the 5,000 probes with the greatest variance in the tumour samples were selected for clustering. The RPMM clustering method is particularly suited to analysis of methylation data generated from the HumanMethylation450 array as output β values fall between 0 and 1, and can be modelled using a β-like distribution (53). For motif analysis, the CentriMo tool was used (19). CentriMo identifies overrepresented motifs within sequences, correlating these with known DNA-protein binding motifs (19). β values were transformed to M values using M=log_2_[ß/(l-ß)]. For differential methylation analysis versus the subset of normal mucosal samples, a probe was considered to be differentially methylated in a comparison if the Benjamini-Hochberg adjusted P value for the comparison was <0.05 and had an average absolute Δß > 0.2 versus normal mucosal samples.

Expression data were preprocessed and normalized using quantile normalization with the Limma R package. For between group comparisons the empirical Bayes function was used, and adjusted for multiple testing using the Benjamini-Hochberg method (55) to control for false discovery rate (FDR) and avoid type 1 errors. We considered 0.05 to be the FDR threshold for significance. For integrated expression and methylation data analysis, genes were considered to be methylated if one probe within 2 kb upstream of the gene transcription Start site (TSS) was differentially methylated by FDR and had an average Aß > 0.2 at that site. If a gene met this criterion, and had a significant FDR corrected P value for the cancer versus normal expression value, it was predicted to be influenced by methylation. Single Sample Gene-Set enrichment analysis was used for between groups comparisons of transcriptomes (16). PANTHER was used to assess enrichment in Reactome pathways and Gene Ontology gene sets (56-58). The CMS classifier package was used to classify cancers into CMS as previously reported (8).

To examine the mutational frequency of epigenetic regulators level 3 somatic variant data was downloaded from the Genome Data Commons portal. Silent variants were discarded and epigenetic regulator genes subset from the EpiFactors Database.

### PRC2 and Methylation overlap analysis

Polycomb occupancy was inferred from SUZ12 CHIP-Seq data from hESC1 cells analysed as part of the ENCODE consortium (59). SUZ12 was chosen as a Surrogate for PRC2 occupancy as previous studies indicate that it is an essential subunit of the PRC2 complex (18, 60). The overlap function within BedTools (61) was used to overlap differentially methylated probes within each cluster versus normal with regions where SUZ12 was bound in hESC1 cells, producing a list of regions where methylation and PRC2 occupancy co-occurred.

### Random forest methylation cluster classifier

The random forest algorithm (randomForest in R) (62) was used to classify the TCGA cohort into methylation clusters. For training, we used the initial cohort of 216 cancer samples with known methylation cluster results. Parameters were tuned to optimize the model (ntree = 5,000, mtry =85). The same 5,000 probes identified in the initial consecutive series were a subset from the supplemental cohort matrix, and subsequently predicted using the model built upon the training set.

### Synthetic Lethality Analysis

Cell line dependency data from Meyers et al (25),. was correlated with colorectal cancer cell line mutation data (24). Synthetic lethal relationships were inferred if a high impact mutation (Truncating mutations or those in splice sites) occurred in one subunit of a molecular complex, and the cell line had relatively higher dependence values on other subunits when compared with cell lines that lacked a mutation. Cell lines were grouped as having a mutation in a specific gene and those not having a mutation, and a Students T-Test performed on dependence values every other subunit within the complex.

### Statistical analysis

For statistical analyses a combination of Software were used, including R and GraphPad Prism 7. Fisher’s exact test was used for hypothesis testing on 2×2 contingencies. Pearson’s chi-squared test was used to compare contingencies > 2×2. Student’s t-test or Wilcoxon rank-sum test was used to compare continuous variables where appropriate. One-way analysis of variance (ANOVA) was used for continuous variable comparisons with > 2 groups.

## Declarations

### Ethics approval and consent to participate

The study protocol was approved by the Royal Brisbane and Women’s Hospital and QIMR Berghofer Medical Research Institute Research Ethics Committees. Written informed consent was obtained from each patient prior to inclusion in the current study.

### Consent for publication

Not applicable

### Availability of data and material

These data has been deposited in the ArrayExpress database at EMBL-EBI (www.ebi.ac.uk/arrayexpress) under the following accession number: E-MTAB-7036 (Methylation)

### Competing interests

The authors declare they have no competing interests

### Funding

This work was supported through funding from the National Health and Medical Research Council (Grant #: 1050455, 1063105), the U.S. National Institutes of Health (Grant #: R01 CA151933 and R35 CA197735), and Pathology Queensland. VW is the recipient of a Senior Research Fellowship from the Gastroenterological Society of Australia. LF was supported by a Research Training Program Living Scholarship from the Australia Government and a Top-Up award from QIMR Berghofer.

Funding agencies that supported this work had no a role in the design of the study, nor did they have input into the collection, analysis or interpretation of data.

### Authors’ contributions

LF performed bioinformatic and statistical analyses on the data, was involved in the conceptualization of aspects of the study and prepared the manuscript.

TD performed molecular and bioinformatic analyses and revised the manuscript for content.

GH was involved in statistical and bioinformatic analysis of the data.

KN was involved in bioinformatic analysis of methylation data and revised the manuscript for content.

CB performed molecular analyses and revised the manuscript for content.

DM was involved in molecular analyses.

LB processed the microarrays.

GM processed the microarrays.

LW was involved in bioinformatic analyses.

KK was involved in bioinformatic analyses.

AMP was involved in bioinformatic analyses of TCGA exome data.

SK was involved in bioinformatic analyses of TCGA exome data.

JP was involved in bioinformatic analyses of TCGA exome data.

NW was involved in bioinformatic analyses of TCGA exome data,

PW performed CMS analysis,

PL performed LINE1 methylation assays and analysis,

YI performed LINE1 methylation assays and analysis,

SO performed LINE1 methylation assays and analysis and provided supervision for this aspect of the study,

RS supervised the study and was involved in conceptualization of aspects of the study,

ST performed CMS analysis and provided supervision for this aspect of the study,

BL supervised the study, was involved in conceptualization of aspects of the study, revised the manuscript for content and secured funding for the study

VW conceptualized the study, performed statistical analyses, revised the manuscript for content, secured funding for the study and provided overarching supervision of the study.

### Acknowledgemens

Not Applicable

### Authors’ information

Not Applicable

